# PIP-Seq identifies novel heterogeneous lung innate lymphocyte population activation after combustion product exposure

**DOI:** 10.1101/2024.06.24.600420

**Authors:** Yung-An Huang, Xinyu Wang, Jong-Chan Kim, Xiang Yao, Anshika Sethi, Allyssa Strohm, Taylor A. Doherty

## Abstract

Innate lymphoid cells (ILCs) are a heterogeneous population that play diverse roles in airway inflammation after exposure to allergens and infections. However, how ILCs respond after exposure to environmental toxins is not well understood. Here we show a novel method for studying the heterogeneity of rare lung ILC populations by magnetic enrichment for lung ILCs followed by particle-templated instant partition sequencing (PIP-seq). Using this method, we were able to identify novel group 1 and group 2 ILC subsets that exist after exposure to both fungal allergen and burn pit-related constituents (BPC) that include dioxin, aromatic hydrocarbon, and particulate matter. Toxin exposure in combination with fungal allergen induced activation of specific ILC1/NK and ILC2 populations as well as promoted neutrophilic lung inflammation. Oxidative stress pathways and downregulation of specific ribosomal protein genes (*Rpl41* and *Rps19*) implicated in anti-inflammatory responses were present after BPC exposure. Increased IFNγ expression and other pro-neutrophilic mediator transcripts were increased in BPC-stimulated lung innate lymphoid cells. Further, the addition of BPC induced *Hspa8* (encodes HSC70) and aryl hydrocarbon transcription factor activity across multiple lung ILC subsets. Overall, using an airway disease model that develops after occupational and environmental exposures, we demonstrate an effective method to better understand heterogenous ILC subset activation.

## Introduction

Since their characterization, group 2 innate lymphoid cells (ILC2s) have been found to play a critical role in the pathogenesis of eosinophilic type 2 inflammation that is present in most forms of human asthma^1-3^. ILC2s are lineage-negative cells that are largely tissue-resident cells and rapidly respond to a multitude of stimuli (allergens, infectious agents, and toxins). ILC2s are a subset of the broader group of innate lymphoid cells (ILCs) and are defined by their production of type 2 cytokines that include IL-5 and IL-13. In contrast, group 1 innate lymphoid cells (ILC1s) produce IFN-γ that are innate counterparts to CD4+ Th1 cells. In mouse models of type 2 asthma, ILC2s play a critical role in the development of type 2 lung inflammation induced by house dust mite^4^, ovalbumin^5^, and *Alternaria alternata*^6^. In addition to allergens, lung ILC2s are stimulated during exposure diesel exhaust, particulate matter, and ozone and contribute to lung inflammation induced by pollutants^7-10^.

The effect of toxic injuries and innate lymphoid cell responses may be relevant to occupational lung diseases such as those that occur in military populations that have been exposed to burn pits and other deployment-related toxins^11^. Burn pits emit airborne toxins during the combustion of debris and leftover material that includes plastics, ordinance, paints, fuel, and oil and are associated with significant health consequences in military personnel and Veterans^12-14^. New-onset and worsening of asthma and chronic obstructive pulmonary disease (COPD) are specifically associated with burn pit exposures^12-20^. There is limited data regarding which burn pit components may induce aberrant airway immune responses, but likely candidates include fine particulate matter (PM), polyaromatic hydrocarbons (PAH), and dioxins^14,20^.

Recent studies have demonstrated that ILC2s are a heterogeneous population with significant plasticity and display regulatory functions as well as produce mediators associated with neutrophilic inflammation^21,22^. Single-cell RNA sequencing (scRNA-seq) is a powerful transcriptomic approach to identify subsets of ILC2s in various disease states and organs, and has demonstrated significant ILC2 heterogeneity and tissue-specific responses in both humans ^23,24^ and mice ^25,26^. One of the limitations of in-depth ILC work is the low cell numbers in tissues (approximately 10,000 to 20,000 in naïve mouse lung). While previous studies have used sorted cells based on lineage negative and CD127+ in human or Th1.2+ in mouse cells to perform subsequent scRNA-seq, here we describe a novel approach where ILCs are enriched using magnetic negative selection before performing particle-templated instant partition sequencing (PIP-seq). PIP-seq is a next-generation scRNA-seq method that has the advantage of rapid, scalable, and microfluidic -free processing that allows for improved detection of heterogeneous populations^27^. Importantly, our method allows for in-depth ILC analysis while concurrently allowing for the assessment of non-ILC lung cell populations. We developed a preclinical model using *Alternaria alternata* to induce type 2 lung inflammation in mice in combination with a burn pit constituent (BPC) mix that includes dioxin, aromatic hydrocarbon, and particulate matter. Using PIP-seq after magnetic innate lymphoid cell (ILC) selection, we identified unique ILC2 subsets that are activated after BPC exposure and a shift from a predominant type 2 airway inflammation to a mixed type 1 and type 2 inflammatory response.

## Results

### Burn pit constituent (BPC) mix in combination with *Alternaria* induces mixed granulocytic lung inflammation

To investigate the effects of the inhalation of airborne fungal allergen together with combustion toxins on the development of pulmonary diseases, a burn pit constituent (BPC) cocktail (2,3,7,8-Tetrachlorodibenzodioxin, TCDD; Benzo[a]pyrene, BaP; particulate matter <4 μm, PM4) was administered during a short-term *Alternaria alternata* (Alt)-induced lung inflammation model^28^ (Figure 1A). The short-term Alt model induces strong type 2 innate lymphoid cell (ILC2)-driven lung inflammation characterized by an increase in type 2 cytokines, eosinophil infiltration, and mucus metaplasia^6,28-33^. The addition of BPC to Alt induced a more severe inflammatory response with an increase in total leukocyte counts in bronchoalveolar lavage (BAL) fluid compared to Alt alone (Figure 1B). BPC added to Alt led to a slightly reduced percentage of BAL eosinophils (Figure 1C) and a concomitant increase in percentage of BAL neutrophils (Figure 1E). However, the total number of BAL eosinophils was similar to Alt alone (Figure 1D) and we observed a synergistic increase in total BAL neutrophils after BPC exposure (Figure 1F). ELISA of BAL supernatants showed an increase in IFN-γ from mice exposed to Alt plus BPC compared to Alt only (Figure 1G). Consistent with similar eosinophil numbers in Alt groups with or without BPC, levels of T2 cytokines IL-4, IL-5, and IL-13 were also similar (Figure 1H-J). Overall, the addition of BPC to Alt induces a synergistic neutrophilic inflammation while maintaining strong eosinophilic inflammation induced by Alt. Thus, the phenotype induced by BPC is a unique model to utilize single-cell approaches to better understand ILC responses during mixed granulocytic inflammation induced by toxins with allergen exposures, and compare with a more conventional T2 model induced by Alt alone.

**Figure 1.**
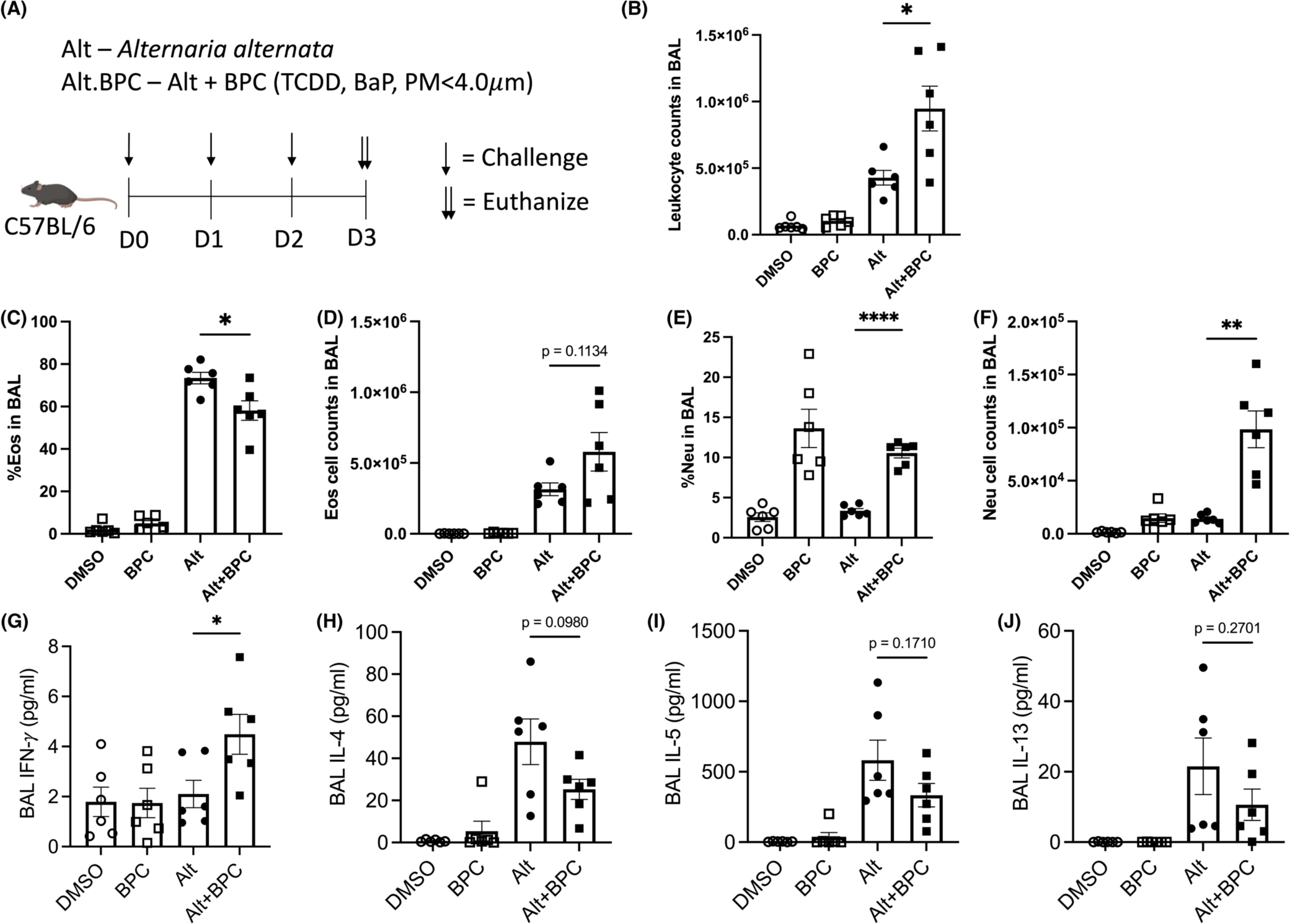
Exposure to burn pit constituents with Alternaria causes a mixed granulocytic lung inflammation. (A) *Alternaria alternata* extract (30 µg/mouse) and burn pit constituent (BPC) cocktail (0.6 ng TCDD + 5 ng BaP + 20 µg PM4/mouse) were administrated intranasally to female wildtype B6 mice on day 0, 1, and 2. The groups are vehicle (DMSO), BPC alone (BPC), *Alternaria alternata* (Alt), and *Alternaria alternata* plus BPC (Alt+BPC). On day 3, mice were euthanized, and bronchoalveolar lavage (BAL) was obtained. (B) Total leukocyte numbers (C and D), leukocyte percentages, eosinophil numbers (Eos), eosinophil percentages, (E and F), neutrophil numbers, and neutrophil (Neu) percentages in BAL were determined by flow cytometry. The concentration of (G) IFN-*γ*, (H) IL-4, (I) IL-5, and (J) IL-13 in BAL was identified by cytokine multiplex assay. Data are presented as mean with SEM (*p < 0.05; **p < 0.01; ****p < 0.0001). TCDD: 2,3,7,8-Tetrachlorodibenzodioxin, BaP: Benzo[a]pyrene, PM4: particulate matter <4 μm

### PIP-seq analysis of enriched lung ILCs reveals unique ILC populations

Given that lung ILCs are a low percentage of the total lymphoid population (less than 1% total lymphocytes), we used magnetic beads to enrich for ILCs before analyzing samples using single-cell RNA sequencing with PIP-seq (Figure 2A). After magnetic separation, ILCs were enriched approximately 100-fold in lung cell samples from both Alt and Alt + BPC samples. Using R package Seurat, we found 19 primary cell clusters (0-18) that were distributed by uniform manifold approximation and projection (UMAP, Figure 2B). To further select for cell clusters containing ILCs, we used the pan-leukocyte marker CD45 (gene symbol: *Ptprc*) as well as CD90 (gene symbol: *Thy1*) (Figure 2C). *Thy1* is primarily expressed by T cell and ILC populations and we next selected 0, 4, 7, 8, and 16 *Th1*-expressing clusters for higher-resolution analysis. There were 13 cell clusters (0-12) identified by sub-clustering (Figure 2D and Figure S1), which were all *Ptprc* and *Thy1* double positive (Figure 2E). We used HSC markers (*Cd34*, *Cd38*, and *Esam*; Figure 2F) and T cell markers (*Cd3e*, *Trac*, and *Trgc*; Figure 2G) to exclude HSC and T cells respectively. The ILC clusters were identified as clusters 0-5, 7, 8, and 11 (Figure 2D).

**Figure 2.**
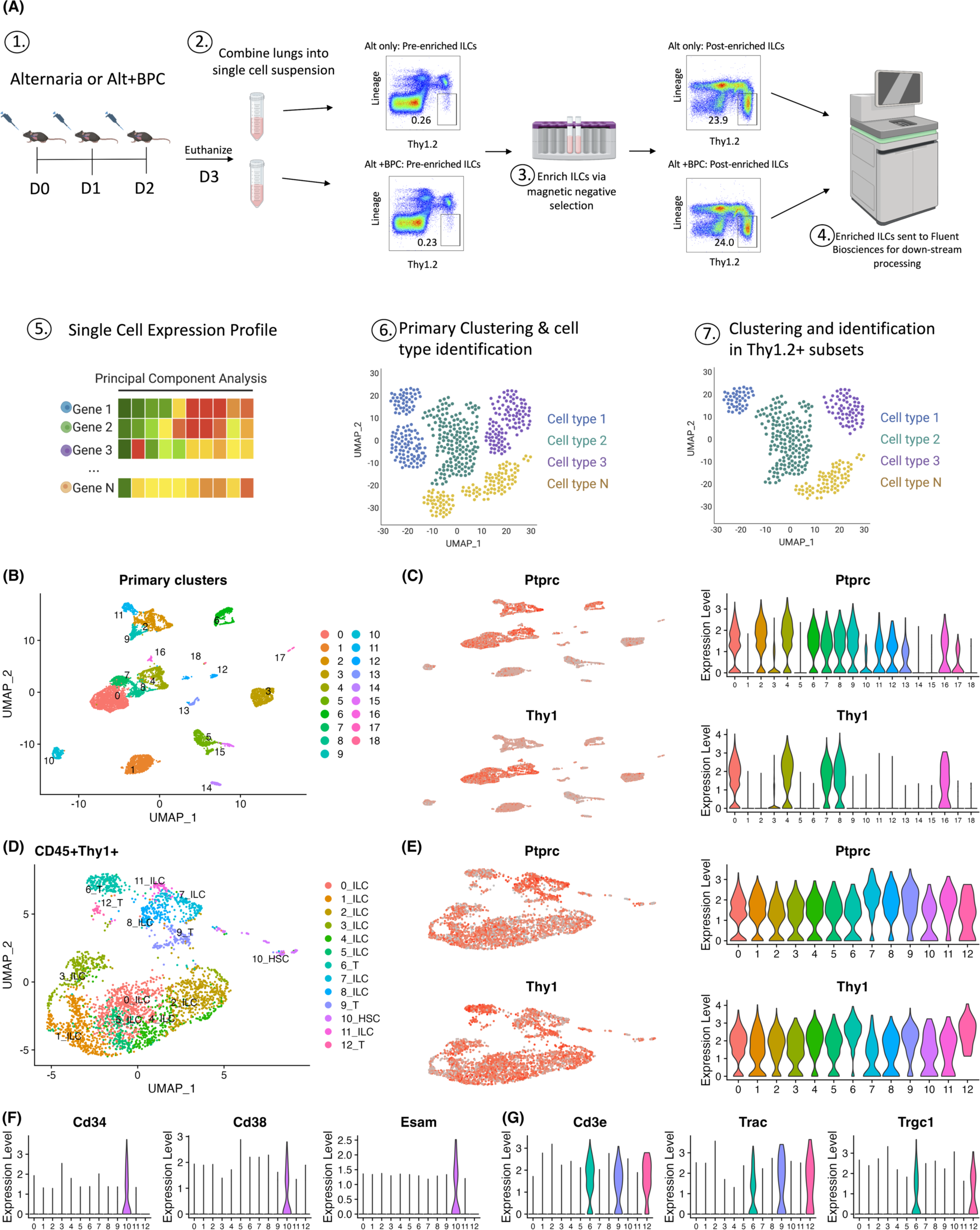
ILC enrichment using magnetic beads improves the ability to identify distinct ILC subsets using single cell RNA-sequencing (scRNA-seq). (A) Preparation of scRNA-seq samples. Female wild type B6 mice were challenged on day 0, 1, and 2 with either Alt only or Alt plus BPC. On day 3, mice were euthanized and lungs were processed into single-cell suspensions after which cells underwent magnetic beads enrichment. Enriched samples were then analyzed using scRNA-seq Single cell expression profiles underwent principal component analysis to generate primary clustering. Clusters that expressed both CD45 (*Ptprc*) and Thy1 (*Thy1*) underwent further subclustering. (B) Distribution of primary cell clusters from enriched samples prior to subclustering. (C) Expression of *Ptprc* and *Thy1* among these clusters. (D) Subclustering based on *Ptprc* and *Thy1* expressing cells. (E) Expression of *Ptprc* and *Thy1* in these subclusters. (F) Expression of hematopoietic stem cell (HSC) markers and T cell markers (G) were identified to differentiate ILCs from T cells and HSCs.

We next identified natural killer (NK) cells and ILCs based on the expression of type 1, type 2, and type 3 signature genes as well as inflammatory cytokine and activation transcription factors, (Figure 3A). ILC clusters 7, 8, and 11 expressed type 1 signature genes (*Tbx1* and *Ifng*). ILC1s (cluster 8) could be distinguished from NK cells (cluster 7 and 11) by the lack of *Prf1*, *Gzma*, and *Gzmb*. Interestingly, cluster 11 also showed expression of type 2 signature genes, including *Il1rl1*, *Klrg1*, *Gata3*, *Rora*, *Arg1*, and *Areg* which suggests they may be non-conventional NK cells, which here we name as ST2 NK. We also found high expression of T2 signature genes in clusters 0-5. We labeled clusters based on peak expression of the following signature genes: cluster 0 (Areg ILC2), cluster 1 (IL-13 ILC2), cluster 2 (IL-10 ILC2), cluster 4 (KLRG1 ILC2), and cluster 5 (LTB ILC2). Cluster 3 had lower expression of effector and activation transcription factor genes compared with other ILC2 populations (but did express *Il13* and *Arg1*) and was labeled as “Quiescent ILC2s”. Surprisingly, we detected minimal type 3 signature genes among all 9 clusters, suggesting that ILC3s were less prevalent compared with ILC2s and ILC1s in our models. We next assessed whether ILC cluster proportions were affected by the addition of BPC to Alt and observed that BPC increased Areg ILC2, LTB ILC2, NK, ILC1, and ST2 NK populations (Figure 3B and C).

**Figure 3.**
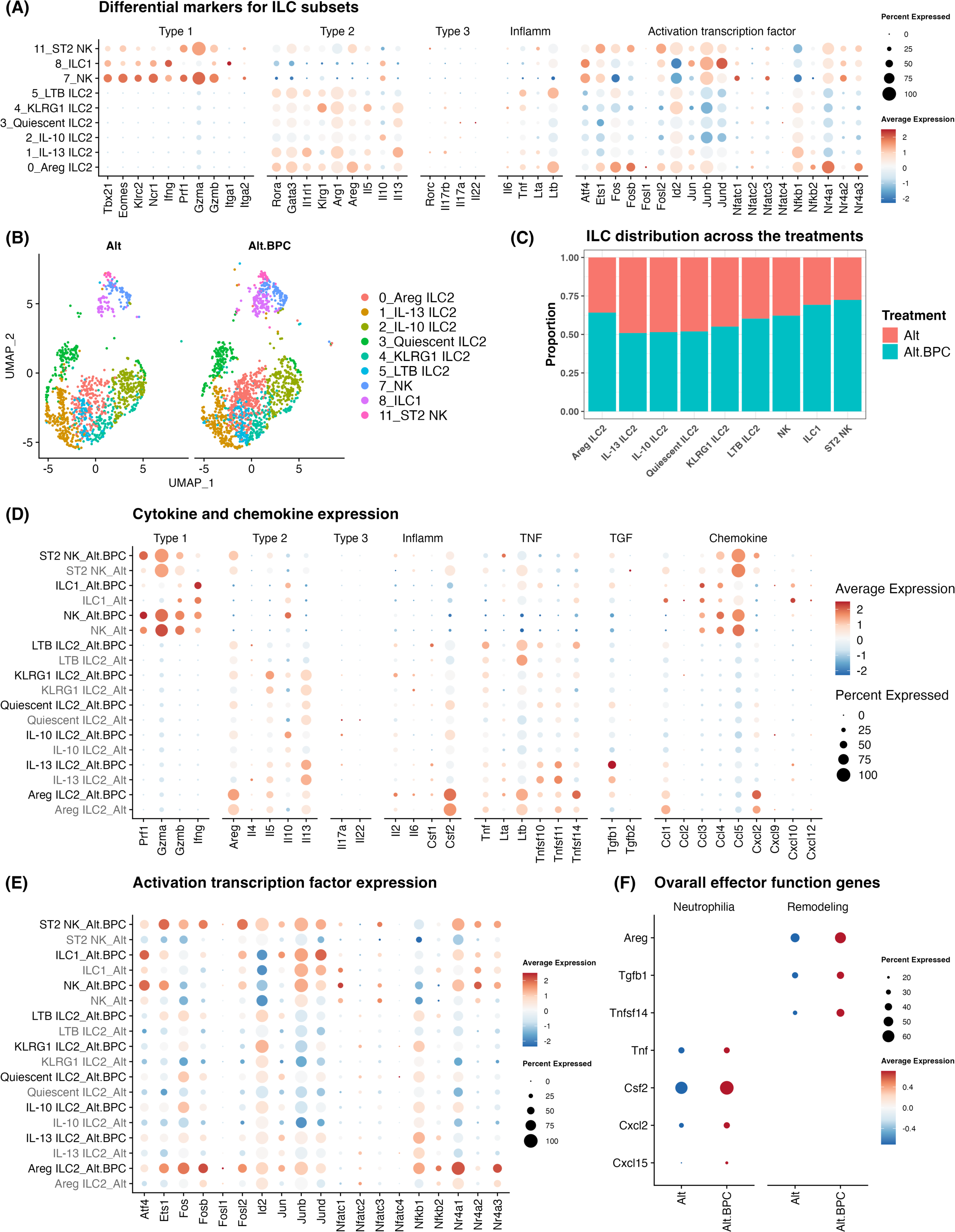
ILC subsets show distinct effector gene expression in response to Alt with BPC. (A) Type 1, type 2, type 3, inflammatory (Inflamm), and activation transcription factor markers were used to identify distinct ILC subsets. (B) The addition of BPC to Alt changes the size of ILC and NK cell clusters. (C) The addition of BPC changes the proportions of ILC subpopulations. (D) Expression of cytokines (type 1, type 2, type 3, inflammatory, TNF super family, and TGF super family) and chemokines in each ILC subset changes after the addition of BPC. (E) The expression of activation transcription factors in ILC subsets changes after the addition of BPC. (F) The overall ILC expression of selected pro-neutrophilic and tissue remodeling-associated effector genes changes after the addition of BPC.

### BPC induces heterogeneous and unique ILC subset activation gene expression

We next assessed expression levels of cytokine, chemokine, and transcription factors within each ILC population after exposure to Alt with and without BPC (Figure 3D and E). Areg ILC2s displayed increased expression of *Areg*, *Ltb*, *Tnfsf14*, *Csf2*, and *Cxcl2* after BPC exposure. Interestingly, though the expression of *Il13* in IL-13 ILC2s increased modestly after BPC, expression of *Tgfb1* was markedly induced in these cells. IL-10 ILC2s showed increased expression of *Il10* after BPC exposure, while *Il5* was highly induced and mainly restricted to the KLRG1 ILC2 subset. In LTB ILC2s, the expression of *Ltb* was decreased after BPC exposure, while the expression of *Tnf* was increased. NK cells showed an increase in *Ifng*, *Prf1,* and *Il10* but reduced *Gzma* and *Gzmb*. ST2 NK cells showed increased expression of *Prf1*, *Gzmb* as well as *Areg*. Robust *Ifng* induction occurred predominately in the ILC1 population, even compared with NK cells, after exposure to BPC. Changes in ILC2 effector mediator expression were consistent with the increase in BAL IFN-*γ* and neutrophilia that we observed at the tissue level with the addition of BPC exposure. Further, levels of *Csf2*, *Cxcl2, Tnf, and Tnfsf14 (encodes* LIGHT) were increased in ILC2s, all of which promote neutrophilic responses. Elevations in *Areg, Tnfsf14* and *Tgfb1* at the cellular levels suggest that BPC may lead to ILC production of tissue remodeling factors (Figure 3F)^34^.

### Unbiased co-expression analysis demonstrates metabolic, IFN-signaling and ribosomal protein transcript changes after BPC exposure

To assess the effect of BPC on lung ILCs in an unbiased manner, we performed co-expression analysis using R package high dimensional weighted gene co-expression network analysis (hdWGCNA). There were 9 gene modules identified and the top 10 hub genes are shown (Figure 4A). We then applied gene set enrichment analysis with WikiPathways database to these modules, and the most relevant pathways are labeled (Figure 4A and B). We identified modules related to NK cell signaling (M1: TYROBP causal network), lymphocyte activation (M2: DNA replication, MAPK signaling pathway (M3), type 2 interferon signaling (M4), cytoplasmic ribosomal proteins (M5 and M9), p53 signaling (M6), *Rora*-related nuclear receptors (M7), and metabolic function (M8: electron transport chain). We then calculated the odds ratio for enrichment of specific modules in type 1 (ILC1 and NK) and type 2 (ILC2) cells between the treatment groups (Figure 4C and 4D). We found that M5 (cytoplasmic ribosomal proteins) and M8 (electron transport chain) were enriched in the Alt-only group. However, the addition of BPC led to the enrichment of M2, M3, and M7 (lymphocyte activation), M6 (p53 signaling), and M1 (TYROBP signaling) among NK, ILC1, and ILC2 cells. Notably, we detected enrichment of M4 (type II IFN signaling) in Alt plus BPC-treated ILC2s, but not in ILC1s and NK cells. This suggests that BPC induces an ILC2 program that includes a response to IFN-*γ* (*Stat1*, *Gpb2*, *Gpb6*) that is produced by ILC1s (Figure 3) and was specifically detected in ILC2s and not found in other ILCs.

**Figure 4.**
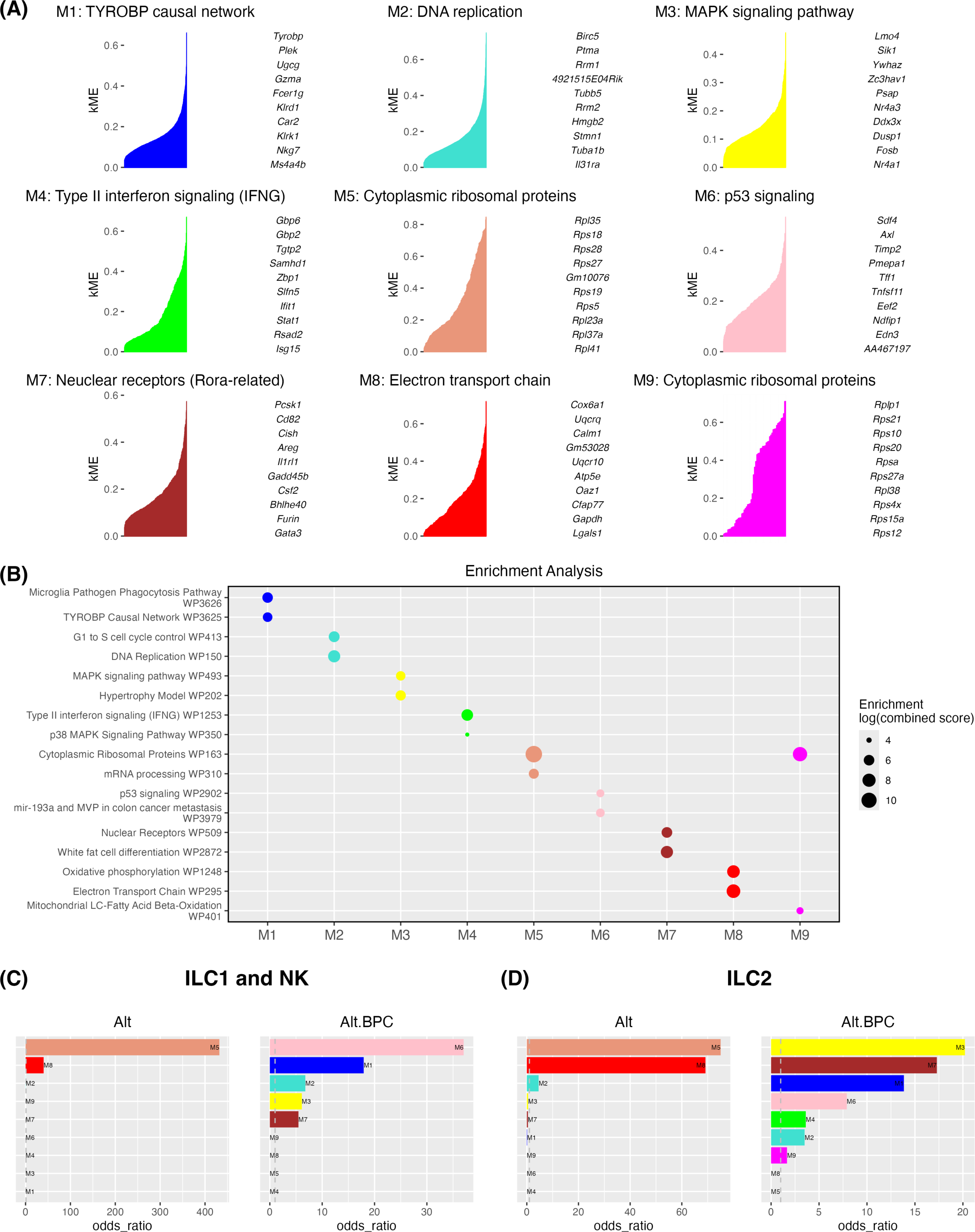
Gene co-expression network analysis reveals metabolic changes and upregulation of activation signaling in ILC subsets after exposure to Alt plus BPC. The co-expression network modules were generated using hdWGCNA (A) The top 10 hub genes with the kME plots are shown with the most relevant biological pathways. The modules underwent pathway enrichment analysis using WikiPathways database. (B) The top 2 potential pathways for each pathway subgroup (M1 – M9) are shown. (C) Odds ratio (statistic to quantify the association between the gene list in the current module and the gene list for the current Term) between ILC1 and NK, and (D) ILC2 between Alt and Alt plus BPC.

We next assessed differences in module expression changes among type 1 innate lymphoid populations, namely ILC1s and NK cells. As might be expected given the role NK cell activation^35,36^, exposure to Alt plus BPC led to the upregulation of TYROBP causal network (M1) in both ILC1s and NK cells (Figure 5A and S2). However, changes in lymphocyte activation (M2), MAPK signaling (M3), P53 signaling (M6) and nuclear receptor group (M7) showed a more heterogeneous response. M2 was increased predominantly in the classical NK cell population and enhanced after BPC exposure suggesting NK cell proliferation induced by BPC. M3 was increased in ILC1 cells, while M6 was decreased. Concomitant activation of MAPK signaling and suppression of p53 pathway could lead to ILC1 survival. M7 (that contains *Areg, IL1rl1, Csf2, Gata3*) and M3 (MAPK signaling) were both increased in ST2 NK cells after BPC exposure which supports that ST2 signaling may be involved in the activation NK cell subpopulation. We did not find a significant change in M4 in NK or ILC1 populations, but we did observe an increase in *Stat1* (downstream of IFN-γ) signaling in ILC1 cells (Figure S2D). M8 was decreased in both ILC1s and NK cells, which suggests BPC may trigger a metabolic change in these cells. Finally, the M5 cytoplasmic ribosomal protein pathway was reduced in all type 1 lymphoid populations though M9 cytoplasmic ribosomal protein pathway was increased only in ILC1s. This suggests that specific ILC ribosomal protein transcript modules are selectively regulated from BPC.

**Figure 5.**
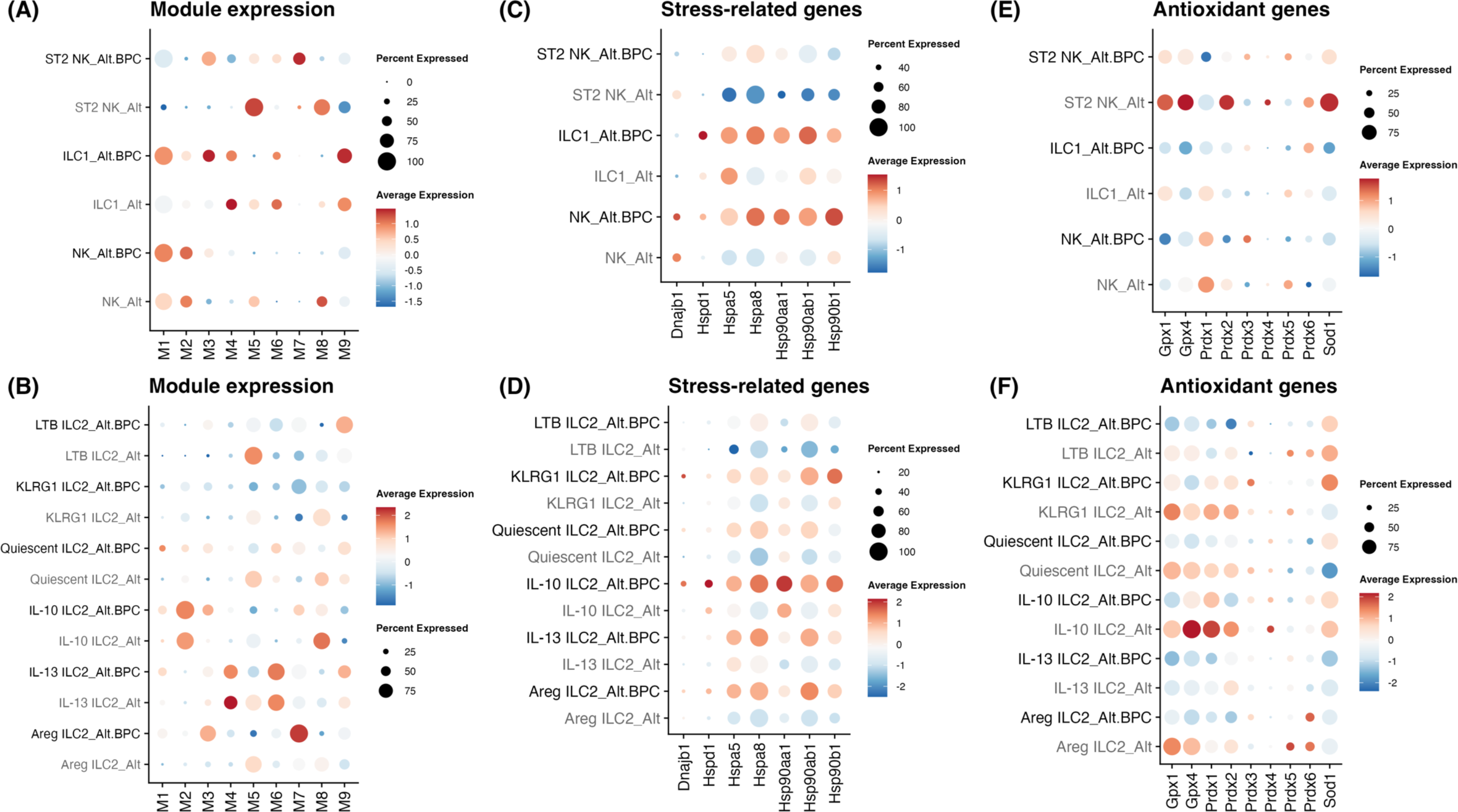
BPC-exposed ILC subsets have increased expression of lymphocyte activation modules and decreased expression of antioxidant genes. The expression of gene co-expression modules in (A) ILC1 and NK, and (B) ILC2 between the treatments are shown in dot plots. Oxidative-stress-related genes in (C) ILC1 and NK, and (D) ILC2 as well as antioxidant genes in (E) ILC1 and NK, and (F) ILC2 between the treatments are also shown in dot plots.

We then applied the same module analysis to ILC2 subpopulations. M1 was not significantly expressed in any ILC2 population (Figure 5B and S3). M2 was significantly expressed in IL-10 ILC2s, and increased after exposure to BPC in these cells, suggesting that BPC may lead to proliferation of IL-10 ILC2s. Type II IFN signaling pathway (M4) was highly expressed in IL-13 ILC2s and showed reduction after exposure to BPC though we also found that *Stat1* was increased after BPC exposure in IL-13 ILC2s (Figure S3D). M3 (MAPK signaling) and M7 (that contains *Areg, IL1rl1, Csf2, Gata3*) were both increased across all ILC2 populations after BPC exposure. M6 was highest in IL-13 ILC2s and further increased after BPC exposure, potentially supportive of increased apoptosis in IL-13 ILC2s. M8 was decreased in all ILC2 populations after BPC exposure, which suggests ILC2s may undergo a metabolic shift from oxidation to glycolysis. Similar to what we observed in type 1 innate cells, the M5 cytoplasmic ribosomal protein pathway decreased in all the ILC2 populations after BPC exposure, but the M9 cytoplasmic ribosomal protein pathway increased. The relatively consistent changes we observed across all innate lymphoid populations in the pattern of M5 and M9 suggest BPC may have a global effect on the expression of ribosomal protein networks in innate lymphoid cells.

Given that air pollutants have been reported to increase oxidative stress, we assessed oxidative stress-related genes and observed increases in the expression of heat shock proteins (HSP) *Hsp90aa1*, *Hsp90ab1*, and *Hspa8* in type 1 and type 2 innate lymphoid populations after BPC exposure (Figure 5C and 5D). Consistent with this, we found a reduction in antioxidant genes *Gpx1*, *Gpx4*, *Prdx1*, *Prdx2*, and *Sod1* in type 1 lymphoid cells (Figure 5E). We also observed a reduction in antioxidant genes *Gpx1*, *Gpx4*, *Prdx1*, and *Prdx2* in ILC2s (Figure 5F). Overall, these results support that BPC exposure promotes ILC oxidative stress (increased oxidant and reduced anti-oxidant gene expression) with overlapping and ILC-specific transcript programs.

### BPC exposure modulates *Hspa8* and select ribosomal gene expression across ILC subsets

We next performed unbiased differential gene expression studies of BPC+Alt versus Alt-treated NK, ILC1, and ILC2 populations. Volcano plots in Figure 6A & B show that *Hspa8* was upregulated in NK cells, ILC1s, and ILC2s suggesting a stress-induced response to BPC in all ILCs. Compared to differential transcripts increased by BPC, there were more highly downregulated transcripts common to all NK, ILC1, and ILC2 populations. Ribosomal protein gene *Rpl41* expression was reduced in all NK, ILC1, and ILC2 subsets. *Rps18* and *Rps19* were reduced in both NK and ILC2 populations. To further study the effect of BPC on innate lymphoid cells, we performed gene set enrichment analysis using gene ontology (GO) database in cellular components on NK, ILC1, and ILC2. The top 10 affected GO terms and the genes associated are displayed in enrichment maps (Figure 6C, 6D). In both type 1 and type 2 ILCs, ribosome- or mitochondria-associated gene components were downregulated with BPC exposure. These data support a consistent effect of BPC on ribosomal protein expression and oxidative phosphorylation in innate lymphoid cells. Enhancement of cytokine binding gene sets occurred in ILC2s after BPC exposure including *Il6ra*, *Il13ra2*, and *Il23r*. Notably, the exposure to BPC plus Alt also led to the reduction of H2 and H4 histone gene expression levels in ILC2s including *H2ac6*, *H2ac7*, *H2ac12*, *H2bc12*, and *H4c17* (Figure 6D). These genes are associated with nucleosome/chromatin formation and support that the addition of BPC may lead to histone-remodeling epigenetic regulation specifically in ILC2s.

**Figure 6.**
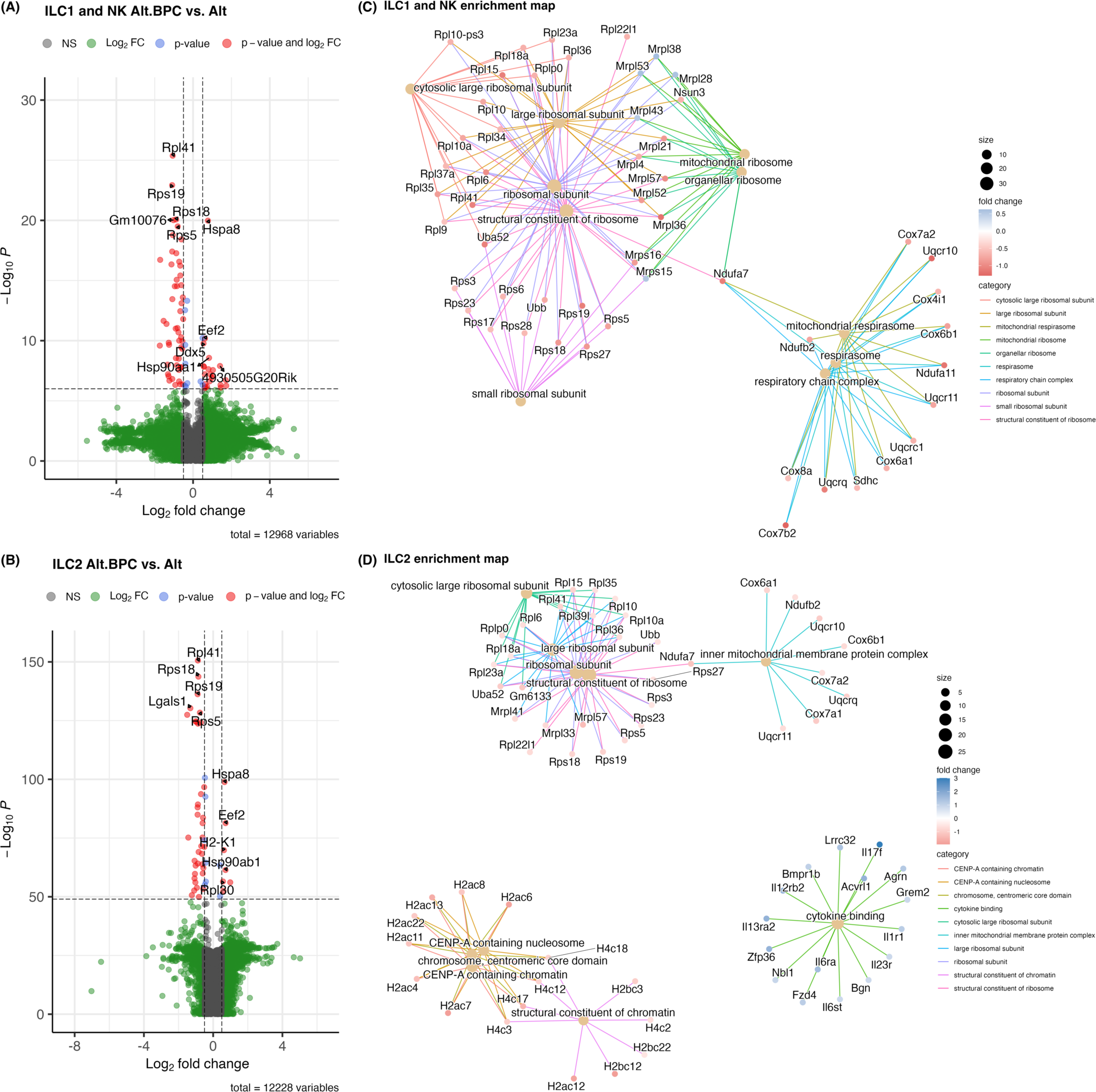
Ribosomal protein and oxidative phosphorylation-related gene expression are reduced after exposure to Alt plus BPC. Most significant variable genes (5 upregulated and 5 downregulated genes) after exposure to Alt plus BPC compared to Alt only are shown for ILC1 and NK (A), and ILC2 cells (B). X-axis is log_2_ fold change while y axis shows -log_10_ *P* value using EnhancedVolcano. The top 10 affected terms that were changed with Alt plus BPC (based on Gene Ontology; GO) are shown for ILC1 and NK (C), and ILC2 (D) cells. Circles represent the category size and colors represent the log_2_ fold change using clusterProfiler and enrichplot.

### BPC exposure transcriptome supports an ILC heat shock protein-aryl hydrocarbon axis

Extracellular heat shock proteins can stimulate immune cells and act as an important form of cellular communication in the lung during stress^37^. To broadly assess HSP-related transcripts from multiple lung cell types, we analyzed CD45^+^Thy1^-^ hematopoietic cell and CD45− structural cell responses after BPC exposure (Figure S4 & S5). Differentially expressed genes were determined in these populations, and the most significant ones (5 upregulated and 5 downregulated) are displayed in volcano plots (Figure 7A & B). Interestingly, as in the ILC populations, *Hspa8* was the most highly expressed transcript in CD45+Thy1− and CD45− cells after BPC+Alt exposure. Upon further analysis of HSP family gene expression, the HSP70 and HSP90 families were elevated in most Thy1− leukocytes and CD45− structural cells (Figure 7C&D). The receptors that bind HSC70 (encoded by *Hspa8*) include LOX-1, SREC-1, FEEL-1, NKG2A, NKG2C, and NKG2D^38^, encoded by *Olr1*, *Scarf1*, *Stab1*, *Klrc1*, *Klrc2*, and *Klrk1*. ILC gene expression of HSP receptors was most highly upregulated in ILC1 and NK cells after exposure to Alt plus BPC (Figure 7E).

**Figure 7.**
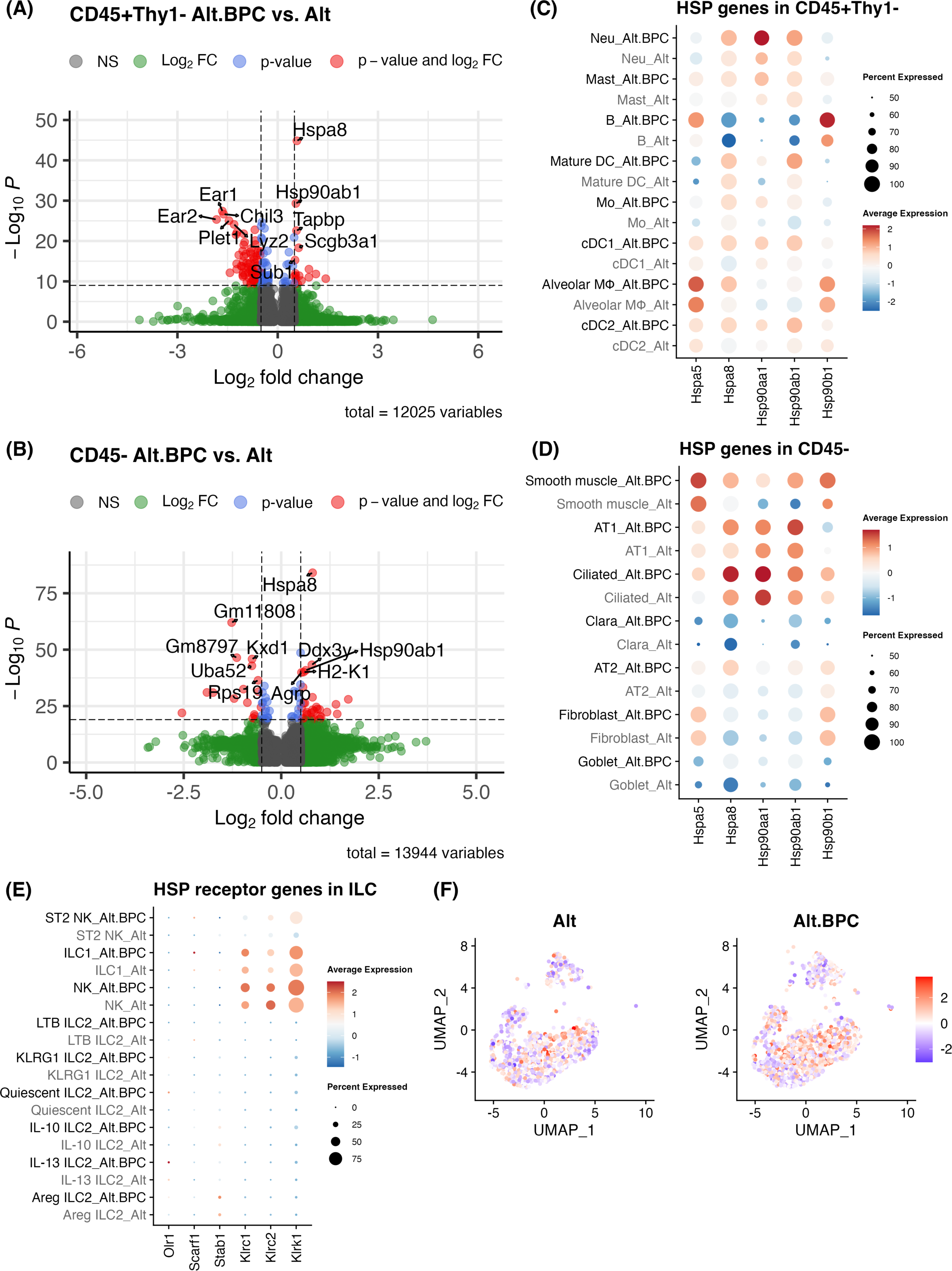
*Hspa8* expression and its downstream signaling is elevated after exposure to Alt plus BPC. Most significant differential genes (5 upregulated and 5 downregulated genes) after exposure to Alt plus BPC compared to Alt only are shown for CD45^+^Thy1^-^ leukocytes (A), and CD45^-^ structural cells (B). X-axis is log_2_ fold change while y axis shows -log_10_ *P* value using EnhancedVolcano. The expression of HSP70 family and HSP90 genes in CD45^+^Thy1^-^ (A), and CD45^-^ (B) between the treatments are shown in dot plots. The expression of HSP70 family and HSP90 receptor genes in ILC (E) between the treatments are shown in dot plots. AHR transcription factor activity is upregulated in multiple ILC populations (F) after Alt plus BPC treatment. Transcription factor activity was determined using decoupleR following Pau Badia-i-Mompel’s workflow, and the inferred activity was plotted in dim plots.

Two of the BPC components, the dioxin TCDD and polyaromatic hydrocarbon BaP have been shown to activate the aryl hydrocarbon receptor (AHR) and lead to oxidative stress^39^. To explore the AHR activity in our model, we assessed the activity distribution of AHR and found greater AHR activity after BPC exposure (Figure 7F). Interestingly, AhR can also be triggered by HSC70^40^ which could further contribute ILC AhR pathway activation after BPC exposure. We assessed inferred AhR transcription factor activity in ILC clusters after BPC exposure and observed enhanced AhR transcription factor activity across multiple ILC clusters (Figure 7F). Thus, there may be direct (TCDD, BaP, PM4.0) and/or indirect (endogenous ligands such as HSC70) activation of ILC subsets through AhR pathways after BPC exposure.

## Discussion

Here we used magnetic bead-based negative selection followed by single-cell RNA sequencing (PIP-Seq) to assess lung innate lymphoid cell transcriptomic changes after mice were exposed to a combination of burn pit constituent (BPC) mix with the fungal allergen *Alternaria*. ILCs are relatively rare populations in tissues which presents challenges for assessing ILC responses amongst other cell types. Our use of ILC enrichment (100x) prior to scRNA-seq differs from prior studies using clustering of sorted cells ILCs^24,41^ ^26^. From our methods, we successfully obtained useful data from non-ILC cell transcriptome analysis (structural and immune cells, Figure 7) while allowing for high-resolution analysis of lung ILC heterogeneity. Another benefit of using bead-based selection to purify for ILCs followed by fluidic-free single-cell sequencing is a reduction in cost and time compared to cell sorting-based selection.

We identified unique ILC subsets in the mouse lung with distinct transcriptomic changes after exposure to BPC with Alt. While Alt induces a well-described type 2 model of eosinophilic lung inflammation^6,33^, the addition of BPC led to a superimposed type 1 inflammation characterized by upregulation of IFN-*γ* and increased airway neutrophilia. Our transcriptomic data demonstrated IFN-*γ* expression in both NK cells and ILC1s. *Csf2* and *Cxcl2* were also upregulated in ILC populations and their products GM-CSF and CXCL2 contribute to tissue neutrophil accumulation. Gene co-expression analysis showed an increase in type II IFN signaling pathways in ILC2s after BPC treatment, but interestingly not in ILC1s or NK cells. This suggests that ILC2s could be responding to IFNγ from ILC1/NK cells. We detected activation of MAPK signaling and suppression of p53 signaling in ILC1s, which suggests BPC may contribute to ILC1 activation and survival pathways. Taken together, these results support that type 1 lung inflammation can be induced in the setting of preserved type 2 inflammation, with upregulation of the former not necessarily reducing the latter. This challenges the classical view that Th1 and Th2 inflammation exists in a see-saw model where blocking one axis may tip the balance in favor of the other. Importantly, studies of mixed granulocytic severe asthma have demonstrated that this endotype has the lowest lung function and more severe disease.^42,43^ Despite this, there is a paucity of models of mixed granulocytic asthma to provide mechanistic insights or targets.

We found upregulation of genes associated with oxidative stress, and reduction in antioxidant genes in both type 1 and type 2 innate lymphoid cells after BPC exposure. Environmental pollutants such as PM2.5 and diesel exhaust particles ^44^ have been reported to induce oxidative stress in immune cells. While high microenvironmental reactive oxygen species have been reported to polarize CD4 T cells towards a Th2 phenotype ^45^ the role of ROS in ILCs is less clear. We also found changes in ribosomal protein expression after BPC exposure. Ribosomal proteins play important roles beyond protein translation and are involved in post-transcriptional gene regulation that has multiple effects on immune cells akin to RNA-binding proteins that regulate ILC2 function^46-51^. We detected a consistent reduction of a few specific ribosomal proteins (*Rpl41* and *Rps19*) in ILC1s and ILC2s after BCP exposure. Interestingly, protein products of *Rpl41* and *Rps19* have demonstrated anti-inflammatory effects in previous reports^46,48,52,53^, and perhaps BPC reduction of these contributes to enhanced ILC activation and inflammation. Additionally, specific histone transcripts were downregulated by BPC in ILC2s. Histone remodeling plays a critical role in modulating gene expression ^54^ and regulating cell proliferation^55^. Further study of these regulatory pathways in ILC2s may provide insights into toxin exposure-mediated lung responses.

The heat shock protein transcript *Hspa8* gene was highly upregulated accross ILC populations after BPC exposure. HSC70 is the protein product of *Hspa8* and a member of the larger HSP70 family. *Hspa8* has been reported to be increased upon exposure to air pollutant^56^ and the extracellular form can trigger the activation of immune cells ^37^. The increase of *Hspa8* was found in almost every lung cell type analyzed in the lung, suggesting that high HSC70 may be present in lungs exposed to BPC. We predict that the aryl hydrocarbon (AhR) receptor may have an important role in our model given that the dioxin TCDD and aromatic hydrocarbon BaP can activate AhR. However, endogenous ligands including HSC70 may also play a role in AhR activation as a previous study showed a regulatory role of HSC70-AhR signaling in a colitis model^40^. Importantly, Ahr could be a target for toxin-mediated lung injury and ILCs are known to be regulated in different ways by AhR^21,39,57^.

In conclusion, our PIP-seq analysis of enriched murine lung ILC populations after Alt and BPC exposure revealed unique diverse populations of both type 1 and type 2 innate lymphoid cell responses with distinct patterns of transcriptional activation. The goal of our studies was to use a novel method to broadly assess lung ILC responses to allergen and toxin exposure that may spur further mechanistic investigations. Our results support a role for ILC oxidative stress, specific ribosomal and heat shock proteins, as well as aryl hydrocarbon receptor signaling as fruitful areas of further study. Overall, our findings have important implications for the impact of environmental pollutants on airway disease that can be driven by ILCs and is relevant to military and other inhalation exposures such as wildfires.

## Materials and Methods

### Burn pit-related constituents

Burn pit-related constituent (BPC) cocktail included 2,3,7,8-Tetrachlorodibenzodioxin, (TCDD; Thermo Fisher Scientific, Waltham, MA), Benzo[a]pyrene (BaP; Millipore Sigma, Burlington, MA), and fine atmospheric particulate matter <4 μm (PM4; National Institute of Standards and Technology, Gaithersburg, MD).

### Mouse model

Female C57BL/6 mice at age of 6-8 were obtained from the Jackson Laboratory (Bar Harbor, ME). To induce the pulmonary inflammation, the mice were intranasally challenged with DMSO (vehicle control), BPC, Alternaria alternata (Alt), or Alt plus BPC for 3 days (Figure 2A). One day after the last challenge, the mice were euthanized by CO2 inhalation followed by secondary organ (lung) removal. The lungs were collected for single cell suspension and ILC enrichment (by Mouse Pan-ILC Enrichment Kit; STEMCELL Technologies, Vancouver, British Columbia, Canada). The dose of Alt was 30 µg/mouse, and the BPC cocktail was mixed by TCDD (0.6 ng/mouse), BaP (5 ng/mouse), and PM4 (20 µg/mouse). All experimental protocols (S10137) were approved by the University of California, San Diego Institutional Animal Care and Use Committee and comply with American Veterinary Medical Association (AVMA) standards. We confirm that all methods were carried out in accordance with relevant guidelines and regulations. We confirm that all methods are reported in accordance with ARRIVE guidelines (https://arriveguidelines.org).

### Bronchoalveolar lavage fluid and lung tissue processing

Female C57BL/6 mice at age of 6-8 were obtained from the Jackson Laboratory (Bar Harbor, ME). To induce the pulmonary inflammation, the mice were intranasally challenged with DMSO (vehicle control), BPC, Alternaria alternata (Alt), or Alt plus BPC for 3 days (Figure 2A). One day after the last challenge, the mice were sacrificed.

After the mouse sacrifice, the tracheas were cannulated, and the bronchoalveolar lavage (BAL) fluid was collected in 0.5+0.6×4 ml 2% BSA (Sigma, St. Louis, MO). The supernatant of the initial 0.5 ml was stored for cytokine detection. The lungs were collected in free RPMI medium and then dissociated for single-cell suspension by the Miltenyi Lung Digest Kit and Dissociator (Miltenyi Biotec, Bergisch Gladbach, Germany). Cells counts were determined by flow cytometer Novocyte (Agilent Technologies, Inc., Santa Clara, CA).

### Flow Cytometry

The staining for flow cytometry was performed on 1×10^6^ BAL or lung cells. Fc receptors were firstly blocked by αCD16/32 (Biolegend, San Diego, CA). For differential granulocyte determination, antibodies against CD45.2 (PerCP-Cy5.5), Siglec-F (PE), GR-1(APC), and CD11c (FITC) were applied. Eosinophils were identified as CD11c^-^Siglec-F^+^, and neutrophils were identified as SiglecF^-^GR-1^+^. For ILC identification, antibodies against CD45.2 (PerCP-Cy5.5), Thy1.2 (eFluor 450), and lineage (lin; FITC). The lin consisted of the BioLegend Lineage cocktail (αCD3e, αLy-6G/Ly-6C, αCD11b, αCD45R/B220, and αTER-119), αCD11c, αNK1.1, αCD5, αFcεR1, αTCRαβ, and αTCRLδ. ILCs were identified as CD45.2^+^lin^-^Thy1.2^+^ lymphocytes.

### Innate lymphoid cell enrichment and single-cell RNA sequencing

ILCs from Alt and Alt plus BPC treated mice were enriched from two single-cell suspended lungs by Mouse Pan-ILC Enrichment Kit (STEMCELL Technologies, Vancouver, British Columbia, Canada). ILC enriched samples were preserved in PIPseq^TM^ T100 3L Single Cell Capture and Lysis Kit and processed by sequencing service (Fluent BioSciences, Watertown, MA).

### Single-cell RNA sequencing analysis and CD45^+^Thy1^+^ cell visualization

The barcode-gene matrices from PIPseeker pipeline were further analyzed with the R package Seurat (v4.3.0)^58^. Following the standard practices to exclude the doublets and low-quality cells, cells that were considered as doublets by R package scDblFinder (v1.15.1)^59^, expressed less than 200 or greater than 4000 genes, or had mitochondrial genes greater than 5% were filtered from the datasets.

Control (Alt) and experimental (Alt.BPC) datasets were undergone Seurat’s workflow of SCtransform normalization and integration to generate the primary clustering at the resolution of 0.3. Uniform manifold approximation and projection (UMAP) was employed for dimensionality reduction and visualization of the data. In the purposes of deeper ILC investigation, the clusters with positive expression of *Ptprc* and *Thy1* were subsetted to undergo a secondary clustering at a higher resolution of 1.0.

### Gene co-expression network and pathway analysis

To identify the gene co-expression modules in ILCs in Alt.BCP, R package high dimensional weighted gene co-expression network analysis (hdWGCNA, v0.3.01)^60^ was applied. The co-expression modules were further undergone the pathway analysis by R package enrichR (v3.2)^61-63^ with the WikiPathways database.

### Gene set enrichment analysis

To identify potential effect from the most variable genes, the differential genes were found by Seurat’s FindMarkers function and visualized by EnhancedVolcano (v1.13.2). The differential genes were further undergone the gene set enrichment analysis using R package clusterProfiler (v4.10.0)^64^ with gene sets from Gene Ontology (GO) cellular component and molecular function. R package enrichplot (v1.22.0) were utilized for visualization of GO enrichment maps.

### Transcription factor activity

To identify the activities of the transcription factors, the inferred transcription factor activity analysis was performed with the R package decoupleR (v2.9.1)^65^ following Pau Badia-i-Mompel’s workflow. R package pheatmap (v1.0.12) was utilized for visualization of the transcription factor activity across the clusters.

## Data availability

The datasets are available in the National Center for Biotechnology Information Gene Expression Omnibus (GEO) repository GSE261844 (https://www.ncbi.nlm.nih.gov/geo/query/acc.cgi?acc=GSE261844).

## Code availability

R scripts are available through github (https://github.com/tdohertylab/Alt.BPC_ILC-enriched-lung_scRNAseq).

## Supporting information

Support methods

## Supplementary figure legends

**Figure S1.**
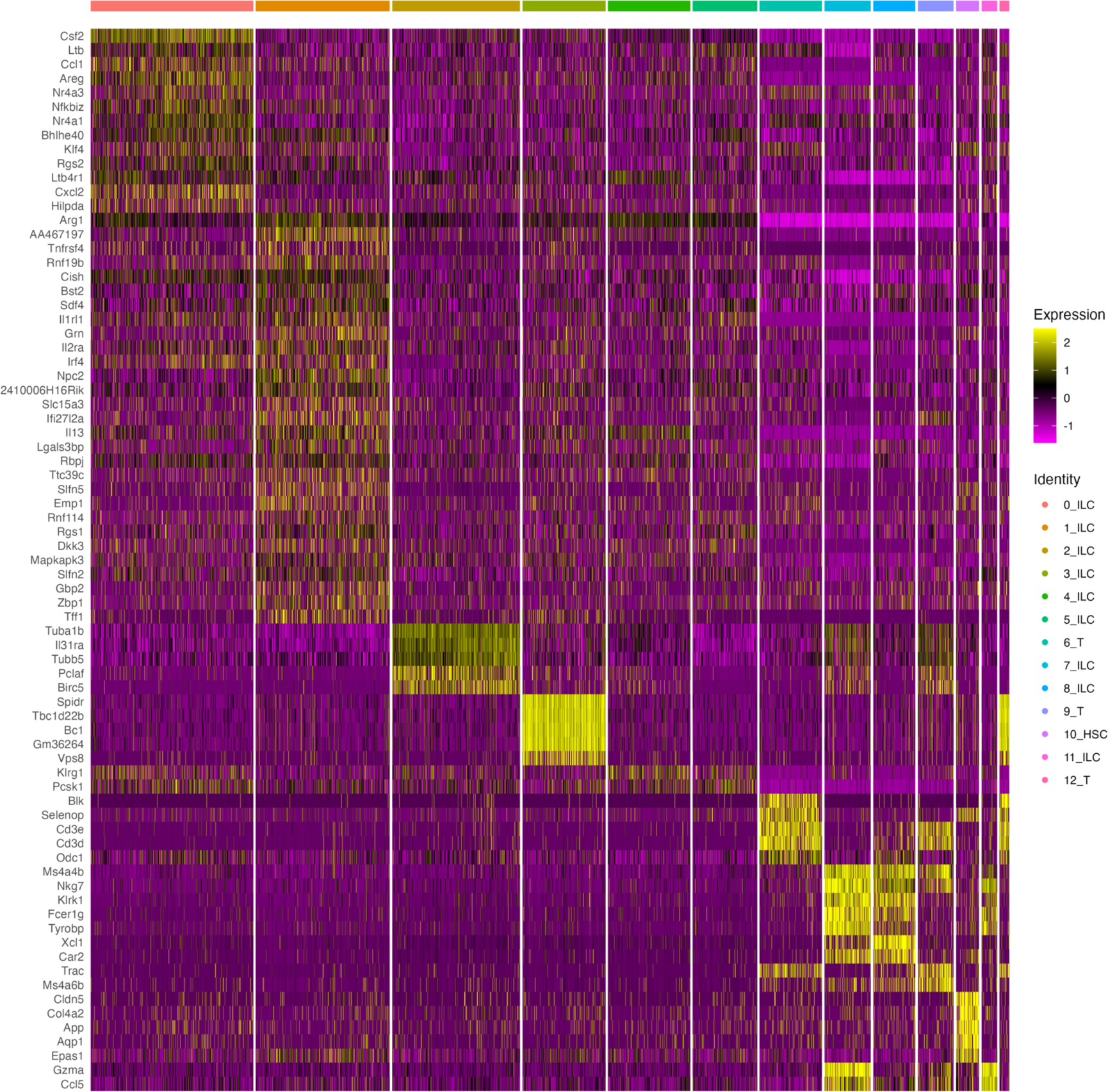
Marker genes for each cluster in CD45^+^Thy1^+^ cell subsets are illustrated as expression level in heat map form.

**Figure S2.**
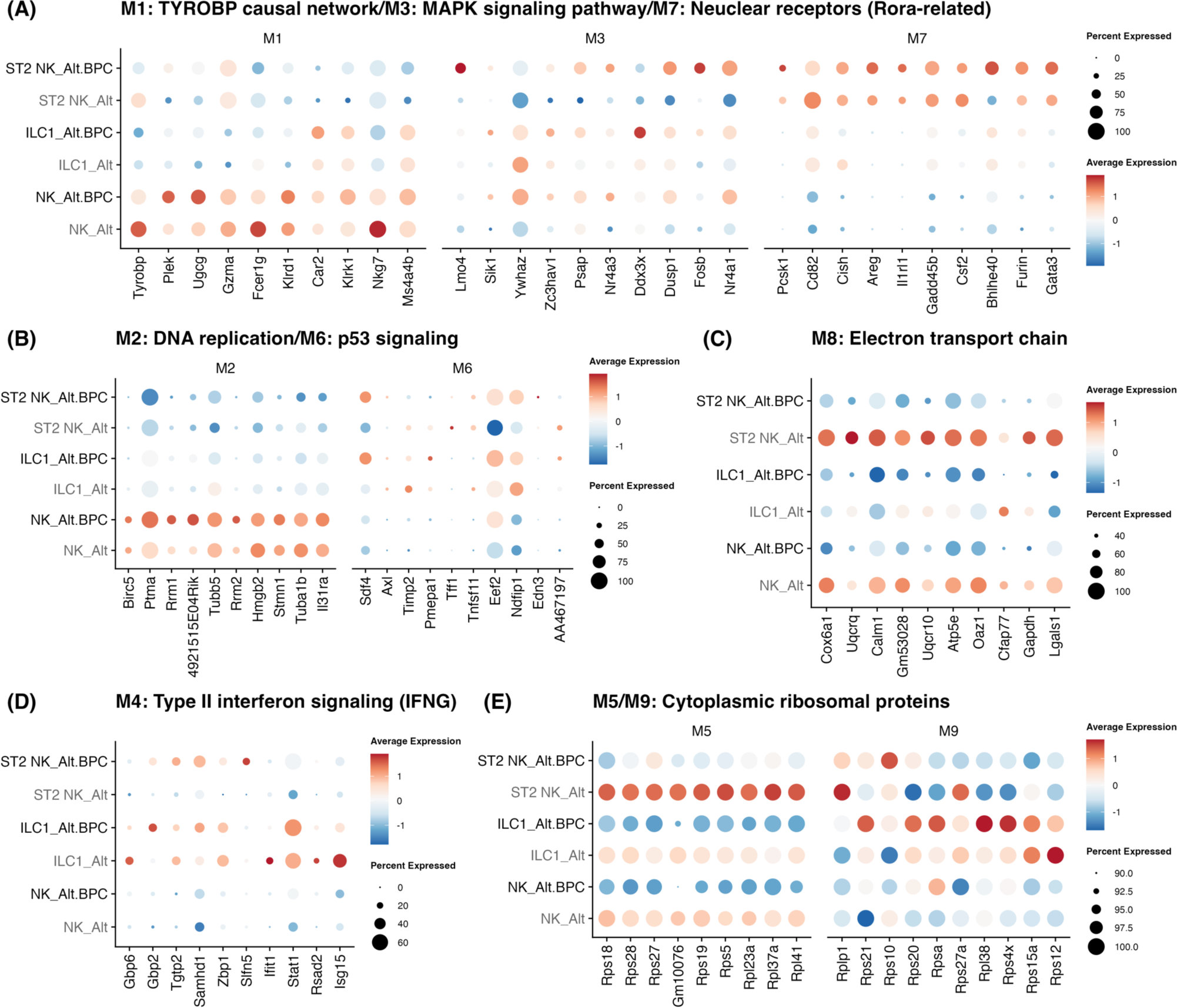
Top 10 hub gene expression from each module in ILC1 and NK are shown in dot plots.

**Figure S3.**
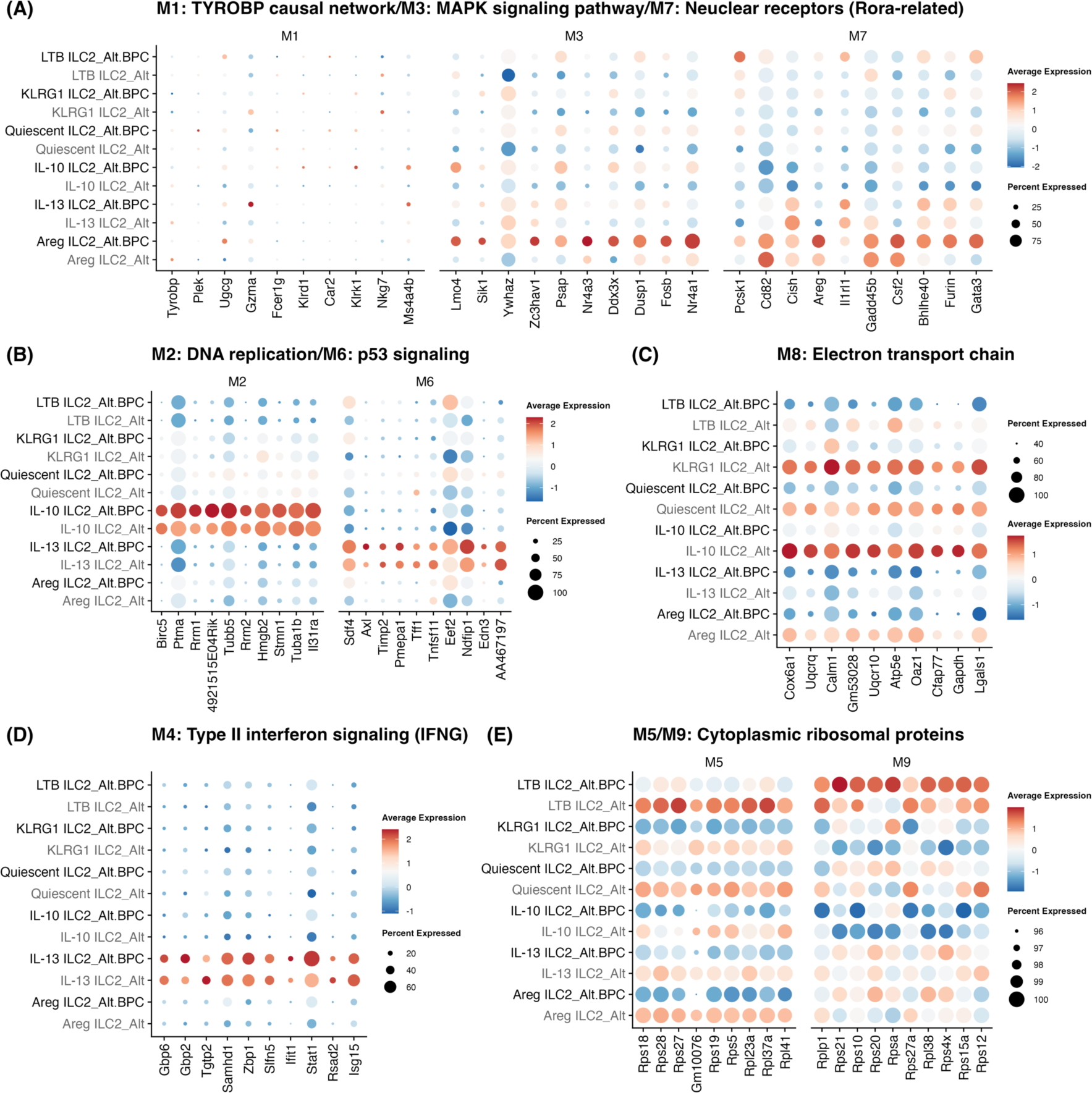
Top 10 hub gene expression from each module in ILC2 are shown in dot plots.

**Figure S4.**
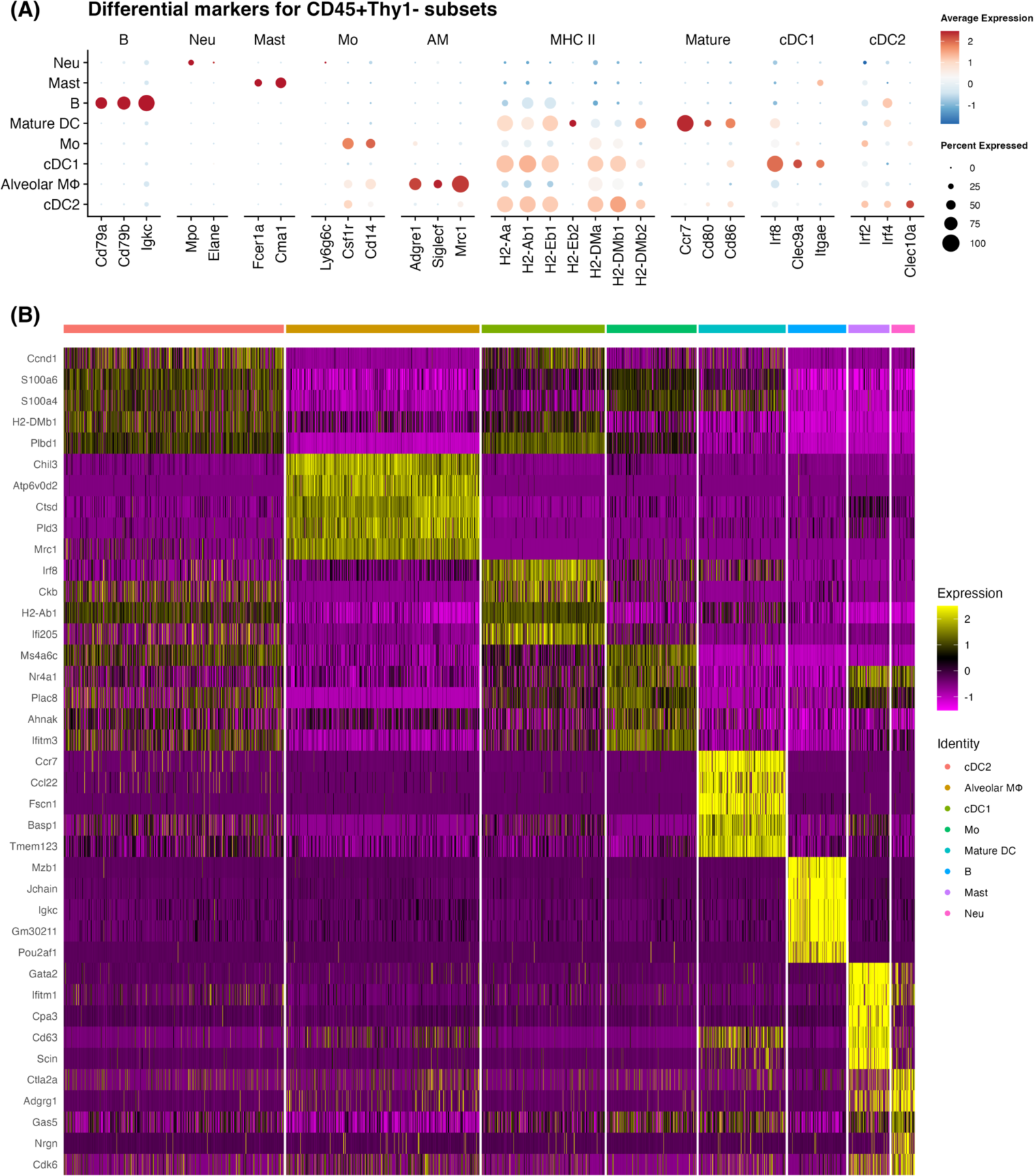
The feature genes (A) for B cell, neutrophil (Neu), Mast cell, monocyte (Mo), alveolar macrophage (AM), MHC class II, type 1 conventional dendritic cell (cDC1), cDC2, mature DC (mature) were used to identify the populations of CD45^+^Thy1^-^ leukocytes. Marker genes for each cluster in CD45^+^Thy1^-^ cell subsets (B) are illustrated as expression level in heat map form.

**Figure S5.**
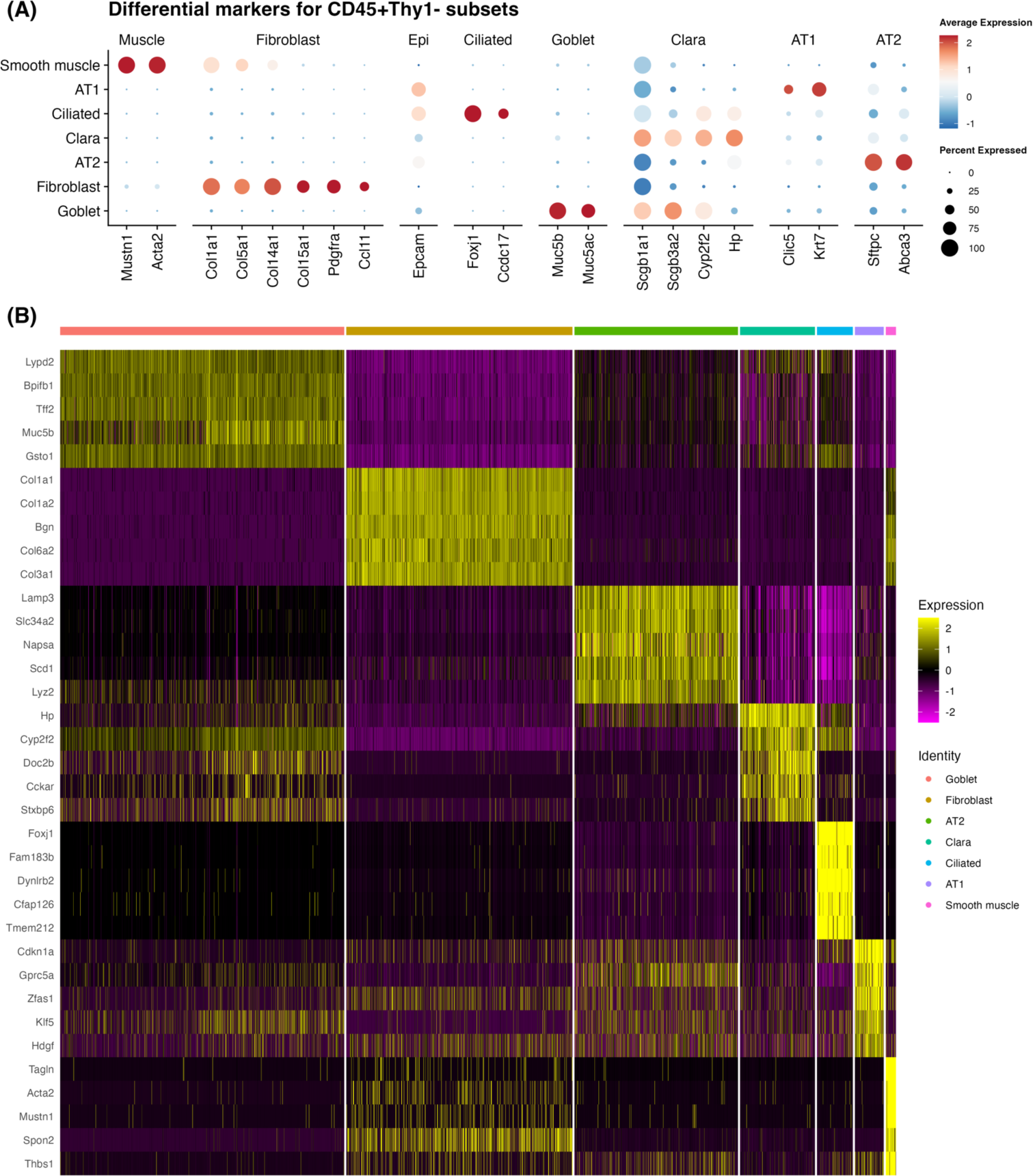
The feature genes (A) for muscle, fibroblast, epithelial (Epi), lung ciliated cell, goblet cell, lung Clara cell, type 1 alveolar cell (AT1), and type 2 alveolar cell (AT2) were used to identify the populations of CD45^-^ cells. Marker genes for each cluster in CD45^-^ cell subsets (B) are illustrated as expression level in heat map form.

