## Supplementary material for "PIP-Seq identifies novel heterogeneous lung innate lymphocyte population activation after combustion product exposure": Support methods

**Supplementary materials and Methods**

**CD45^+^Thy1^-^ and CD45^-^ cell visualization**

After the primary clustering, the clusters with positive expression of *Ptprc* but negative expression of *Thy1* were subsetted to undergo a secondary clustering at the resolution of 0.25. On the other hand, the clusters with negative expression of *Ptprc* were subsetted to undergo a secondary clustering at the resolution of 0.25.
